# A single-cell transcriptomic atlas of T cells across murine lymphoid, mucosal, and gland tissues

**DOI:** 10.64898/2026.11.18.767759

**Authors:** Xiaojing Wang, Biling Pei, Junshuang Liu, Zixuan Han, Xueqing Wang, Yuhan Wang, Xiang Zhao

## Abstract

T cells play essential roles in immune surveillance, host defense, and tissue homeostasis, with their cellular composition and transcriptional states varying across tissues. In this study, we constructed an integrated single-cell transcriptional atlas of 124,186 T cells from seven organs of healthy mice. We identified shared T cell states with distinct tissue associated distributions and transcriptional features. Further analyses revealed transcriptional changes associated with thymocyte maturation, diverse innate-like and memory-like T cell populations in bone marrow, and distinct regional distributions of CD8αα-like T cell states in the intestine. The submandibular gland contained diverse naïve, helper-associated, and effector-associated T cell states. pySCENIC analysis further revealed tissue associated variation in inferred transcription factor regulon activity within shared T cell populations. Together, these findings provide a single cell reference for understanding T cell heterogeneity across lymphoid and non-lymphoid tissues under physiological conditions.

## Introduction

T cells are essential components of the adaptive immune system and play critical roles in host defense, immune surveillance, and tissue homeostasis^1–5^. Their development, maintenance, and immune functions are closely associated with the tissues in which they reside or circulate. The thymus is the primary site of T cell development and selection, whereas the spleen and lymph nodes are major sites for T cell activation and the initiation of adaptive immune responses^6, 7^. Beyond these classical lymphoid organs, other tissues provide distinct environments for T cell maintenance and local immunity. The bone marrow serves as an important niche for memory T cell maintenance^8–10^. The intestinal mucosa harbors diverse conventional and unconventional T cell populations that contribute to epithelial defense and mucosal immune surveillance^11, 12^, whereas the submandibular gland serves as an important effector site for oral mucosal immunity, supporting distinct local adaptive immune responses and tissue-specific immune regulation^13–15^. Together, these organs represent complementary settings for investigating T cell diversity under physiological conditions.

Recent advances in single-cell RNA sequencing (scRNA-seq) have enabled detailed characterization of cellular diversity across tissues. Large-scale studies, including Tabula Muris^16^ and Tabula Sapien^17^, have established reference atlases across mouse and human tissues. Cross-tissue immune profiling has further revealed differences in immune cell composition and transcriptional states among human organs^18^. More recently, Tabula Sapiens 2.0 expanded the human tissue atlas and characterized cell-type-associated transcription factor expression patterns^19^. The CD8 immgenT framework also identified recurrent transcriptional states of mouse CD8αβ T cells across diverse tissues and immune conditions, highlighting their context-dependent distributions^20^. Together, these studies demonstrate the value of cross-tissue single-cell analysis for identifying shared cellular states and tissue-associated differences. Focused characterization of T cells across selected organs may further complement these reference atlases by resolving their cellular and molecular heterogeneity.

In this study, we performed an integrated single-cell transcriptomic analysis of T cells from seven organs of healthy mice: the thymus, bone marrow, lymph nodes, spleen, small intestine, colon, and submandibular gland. We constructed a cross-tissue T cell atlas and examined thymocyte maturation associated states, T cell heterogeneity in bone marrow and secondary lymphoid organs, and regional differences in intestinal T cell populations. Paired single-cell transcriptomic and T cell receptor (TCR) sequencing was additionally used to characterize submandibular gland T cells. Together, these analyses provide a single-cell reference for shared and tissue-associated T cell heterogeneity in healthy mice.

## Result

### A single-cell atlas reveals shared T cell states and tissue-associated compositional differences

To characterize T cell heterogeneity across healthy mouse tissues, we performed single-cell RNA sequencing of CD3⁺ T cells from the thymus, bone marrow, lymph nodes, spleen, small intestine, colon, and submandibular gland (Fig. 1A). For each tissue-derived sample, cells from ten mice (five males and five females) were pooled. After quality control, doublet removal, and data integration, 124,186 T cells from 13 samples were retained for downstream analysis. Unsupervised clustering identified 21 transcriptionally distinct populations, including naive, effector, regulatory, unconventional, cycling, and immature T cell states (Fig. 1B). These populations included several distinct T cell lineages and states, such as CD8αα-like T cells, γδ T cells, NKT-like T cells, and five populations of immature CD4⁺CD8⁺ double-positive (DP) thymocytes. Their identities were supported by the expression of representative lineage and state associated marker genes (Fig. 1E, Supplementary Fig. S1A).

**Figure 1.**
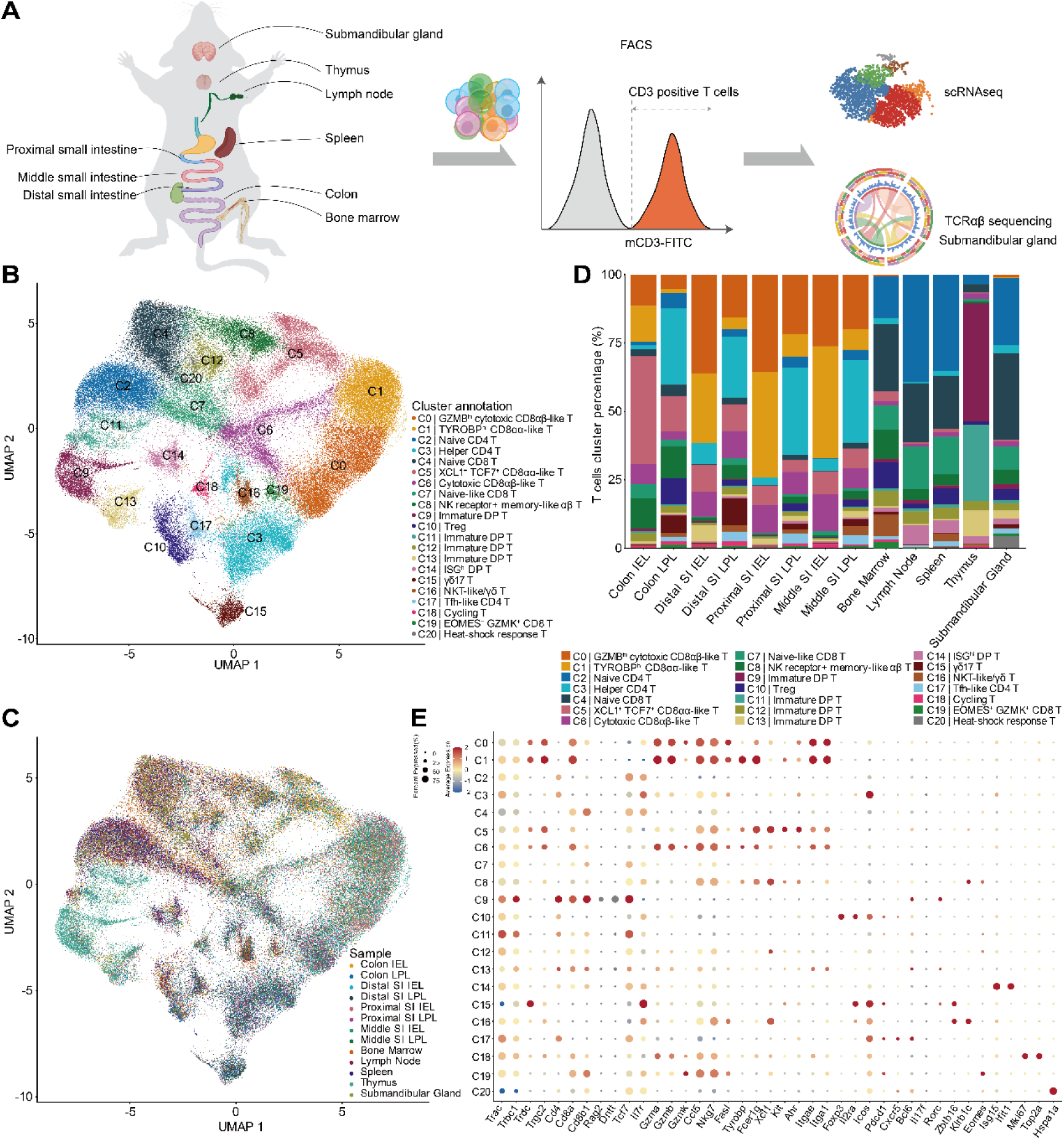
Single-cell landscape of T cells across multiple organs in healthy mice. (A) Schematic overview of the study design and tissue sampling strategy. (B) UMAP visualization of 124,186 T cells collected from 13 samples across seven organs. Each dot represents an individual cell, and colors indicate the 21 transcriptionally distinct T cell clusters. The clusters were annotated as follows: C0, GZMB-high cytotoxic CD8αβ-like TCRαβ/γδ T cells; C1, TYROBP⁺ cytotoxic CD8αα-like TCRαβ/γδ T cells; C2, naive CD4 T cells; C3, helper CD4 T cells; C4, naive CD8 T cells; C5, XCL1⁺TCF7⁺ innate-like CD8αα-like TCRαβ/γδ T cells; C6, cytotoxic CD8αβ-like TCRαβ/γδ T cells; C7, naive-like CD8 T cells; C8, NK receptor⁺ memory-like αβ T cells; C9, immature DP thymocytes; C10, regulatory T cells; C11–C13, immature DP thymocytes; C14, ISG-high DP thymocytes; C15, γδ17 T cells; C16, NKT-like/γδ T cells; C17, Tfh-like CD4 T cells; C18, cycling T cells; C19, EOMES⁺GZMK⁺ CD8 T cells; and C20, heat-shock response T cells. (C) UMAP visualization of T cells colored by tissue region of origin. Each color represents a distinct sampled region. (D) Relative proportions of the annotated T cell clusters across the sampled tissues. (E) Dot plot showing the expression patterns of canonical marker genes used to characterize the identified T cell subsets. Dot size indicates the percentage of cells expressing each gene, and color intensity represents the average expression level.

The integrated atlas revealed the differences in T cell composition across tissues (Fig. 1C, D). The thymus was dominated by immature thymocyte populations, particularly DP T cells, whereas intestinal samples contained diverse transcriptionally distinct CD8αα-like T cell populations. In contrast, lymph nodes, spleen, bone marrow, and submandibular gland shared several major T cell populations but differed substantially in their relative abundance. Naïve CD4⁺ and CD8⁺ T cells were abundant in lymph nodes, spleen, and submandibular gland. In contrast, bone marrow contained a higher proportion of innate-like and effector-associated T cells.

Despite these compositional differences, multiple T cell states were shared across tissues, including naïve, regulatory, and NKT-like T cell populations (Fig. 1E, Supplementary Fig. S1A). pySCENIC analysis further revealed distinct patterns of inferred regulon activity across T cell populations. Some regulons, including Churc1, Fos, and Klf6 associated regulons, showed population-associated differences in mean AUCell scores, whereas Lef1 and Ikzf3 associated regulons exhibited broadly distributed AUCell signals across multiple T cell populations (Supplementary Fig. S1B). Together, these findings reveal tissue associated heterogeneity in T cell composition, shared T cell states across organs, and diverse patterns of inferred regulon activity across T cell populations.

### Transcriptional heterogeneity and inferred maturation trajectories of thymic T cells

To further characterize T cell heterogeneity in the thymus, we reclustered thymus-derived T cells. Distinct CD4⁺CD8⁺ double-positive (DP) and single-positive (SP) thymocyte populations were identified (Fig. 2A). The DP populations included early, intermediate, proliferating, and non-proliferating states. Two DP populations showed relatively high CD4 or CD8 expression, together with elevated expression of selection-associated genes, including *Cd5*, *Cd69*, *Egr1*, *Egr2*, and *Nr4a1*. Different SP populations were distinguished by the expression of representative marker genes (Fig. 2B, Supplementary Fig. S2B). Mature CD4 SP thymocytes expressed maturation and migration associated genes, including *Klf2*, *Sell*, and *Ccr7*. In contrast, the immature CD8 SP population expressed *Runx3* and *Nkg7*. Another distinct CD4 SP population showed elevated expression of *Tnfrsf9* and *Tnfrsf4*. This population also expressed selection associated genes, including *Cd5*, *Nr4a1*, and *Nr4a3*.

**Figure 2.**
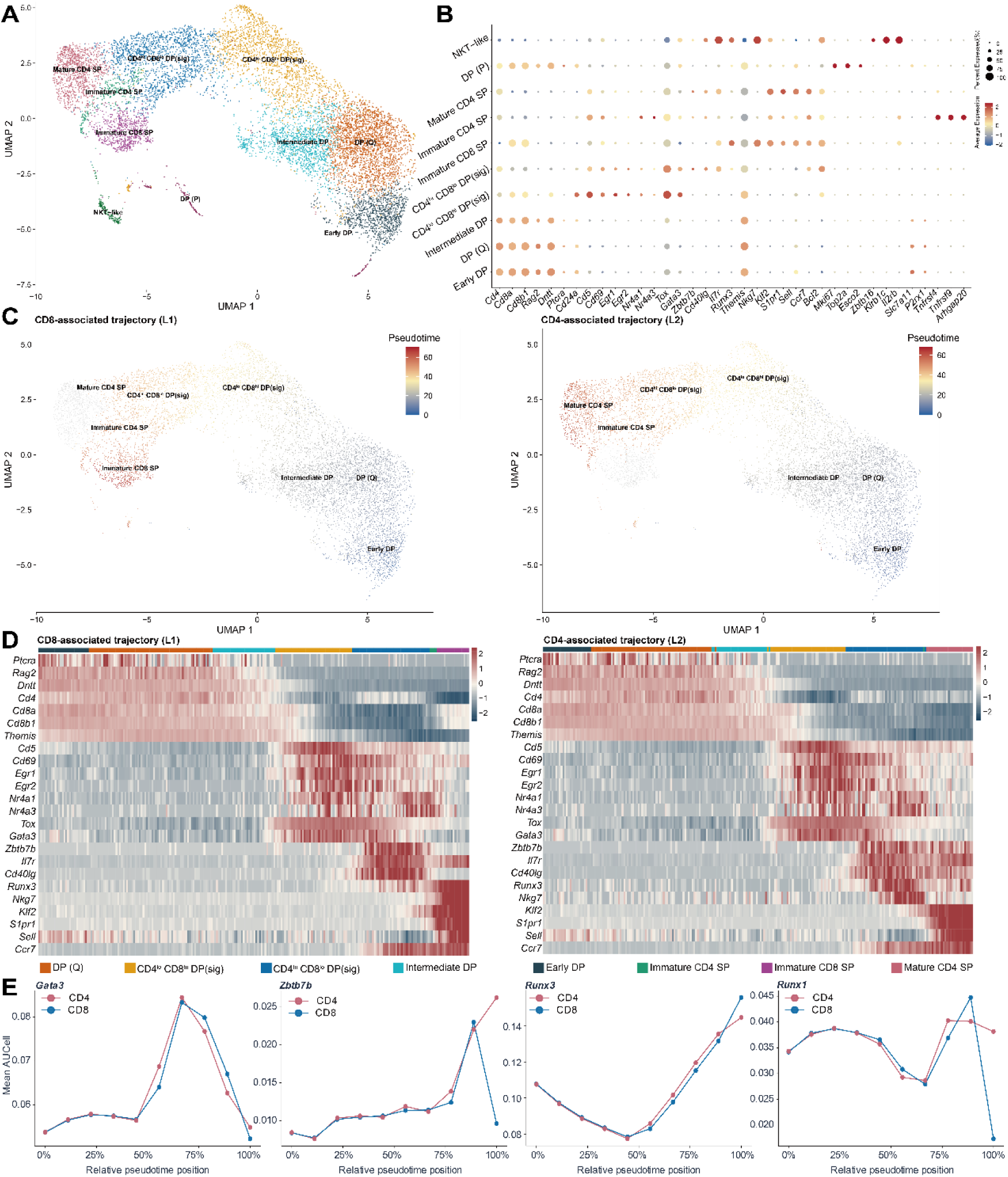
Transcriptional heterogeneity and inferred maturation trajectories of thymic T cells. (A) UMAP visualization of thymus-derived T cells showing ten transcriptionally distinct clusters: Early DP, DP (Q), Intermediate DP, CD4^lo^ CD8^hi^ DP(sig), CD4^hi^ CD8^lo^ DP(sig), Immature CD8 SP, Immature CD4 SP, Mature CD4 SP, DP (P), and NKT-like. Each dot represents an individual cell, and colors indicate the annotated T cell populations. DP, double-positive; SP, single-positive; Q, quiescent; P, proliferating; sig, signaling; lo, relatively low expression; hi, relatively high expression; NKT, natural killer T. (B) Dot plot showing the expression of representative marker genes across the annotated thymic T cell populations. Dot size indicates the percentage of cells expressing each gene, and color represents the average expression level. (C) UMAP visualization of Slingshot-inferred pseudotime trajectories of thymic T cells, colored by pseudotime. Two trajectories are shown: the CD8-associated trajectory (L1) and the CD4-associated trajectory (L2). (D) Heatmap showing the dynamic expression patterns of selected genes along the inferred maturation trajectories, ordered by pseudotime. (E) Line plots showing the mean AUCell scores of four selected transcription factor regulons inferred using pySCENIC across ten pseudotime bins.

We next used pySCENIC to examine inferred transcription factor regulon activity across thymocyte populations (Supplementary Fig. S2B). Tcf7, Tcf12, and Bcl11b associated regulons showed higher z scores in early DP thymocytes. Gata3, Egr1, and Ikzf1 associated regulons showed increased scores in DP populations with selection-associated transcriptional features. Several SP populations showed similar activity patterns for Runx3, Foxo1, and Bach2 associated regulons. In addition, NKT-like T cells exhibited distinct activity patterns for Tbx21, Maf, and Sox13 associated regulons. These results revealed distinct patterns of inferred regulon activity across thymocyte developmental states.

To investigate transcriptional changes during thymocyte maturation, we performed pseudotime analysis using Slingshot. Early DP thymocytes were selected as the starting population. NKT-like cells and the small proliferating DP population were excluded from the analysis. Slingshot inferred two trajectories associated with CD4 SP and CD8 SP thymocyte states. The CD4 and CD8 associated trajectories shared a series of transcriptional states before diverging into distinct lineage associated states. (Fig. 2C and Supplementary Fig. S2A). *Ptcra*, *Rag2*, *Dntt*, and *Themis* were expressed during the early stages of thymocyte development. As differentiation progressed, selection associated genes, including *Cd5*, *Cd69*, *Egr1*, *Egr2*, *Nr4a1*, and *Tox*, showed increased expression. At later stages, the expression of maturation and migration associated genes, including *Klf2*, *S1pr1*, *Sell*, and *Ccr7*, increased (Fig. 2D).

We next examined lineage associated transcription factor expression and regulon activity along the inferred CD4 and CD8 associated trajectories (Fig. 2D,E). Along the CD8 associated trajectory, *Gata3* and *Zbtb7b* expression increased at intermediate stages, followed by a decline in *Zbtb7b* and increased *Runx3* expression toward the terminal stage. In contrast, *Zbtb7b* expression remained at high level along the CD4 associated trajectory. The corresponding regulons also showed dynamic AUCell pattern. Notably, Runx1 associated regulon scores declined at the terminal stage of the CD8 associated trajectory but remained elevated along the late CD4 associated trajectory. These findings suggest that the inferred CD8 associated trajectory transiently exhibits CD4 lineage associated transcriptional features before developing a more prominent CD8 associated program^21^.

### Distinct T cell compositions and inferred regulatory features across bone marrow, lymph nodes, and spleen

To further characterize T cell heterogeneity in bone marrow (BM), lymph nodes (LN), and spleen, we reclustered T cells from these three tissues. Distinct T cell populations were identified, including naïve, memory-like, effector, regulatory, and innate-like T cell states (Fig. 3A). These populations were distinguished by the expression of representative marker genes (Fig. 3B, Supplementary Fig. S3A).

**Figure 3.**
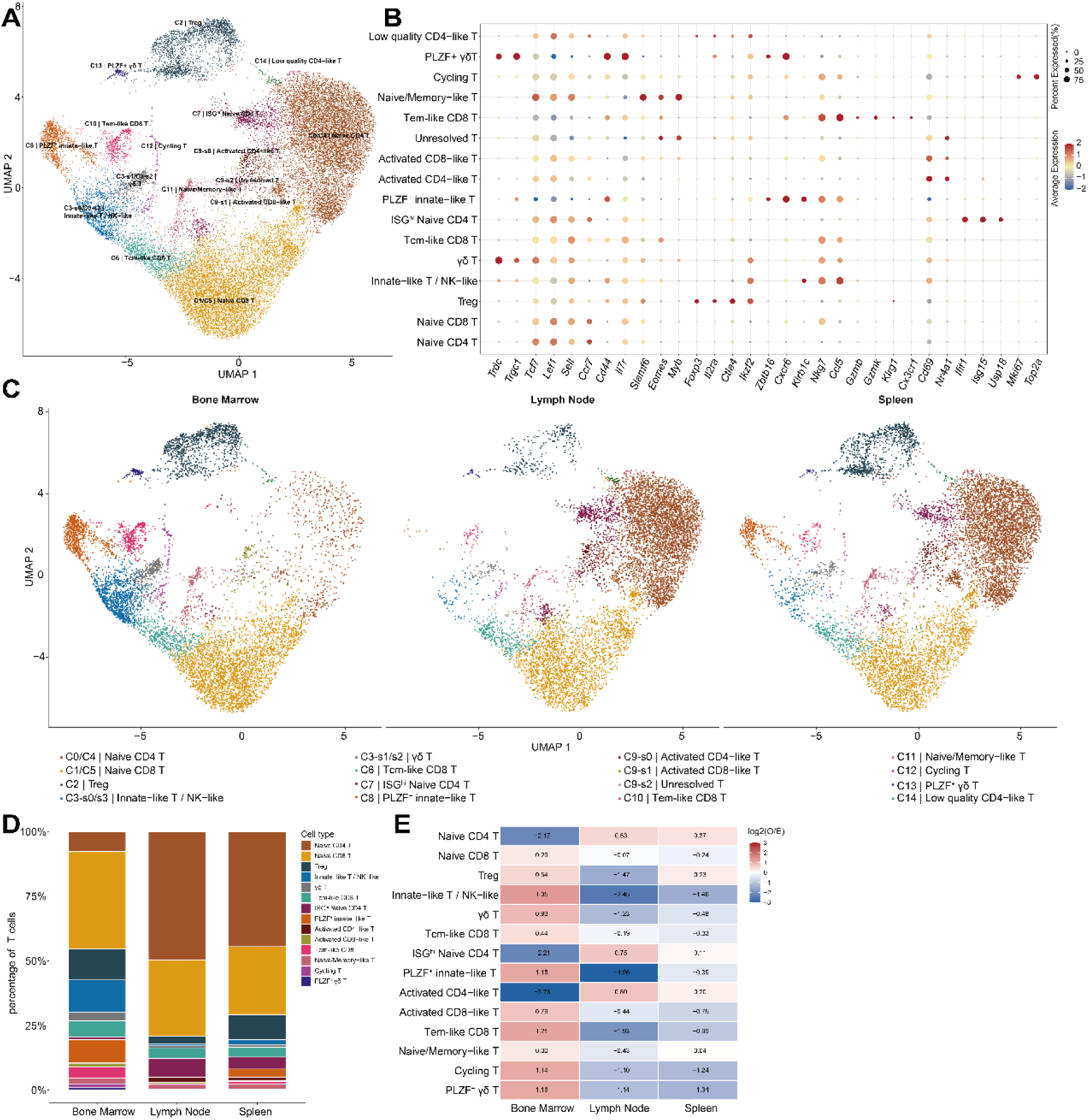
Single-cell transcriptomic characterization and tissue-associated distribution of T cell populations in bone marrow, lymph nodes, and spleen. (A) Uniform manifold approximation and projection (UMAP) visualization of T cells from bone marrow (BM), lymph nodes (LN), and spleen, using the original UMAP embedding with cell annotations refined through subsequent subclustering. Each dot represents an individual cell, and colors indicate the annotated T cell populations. The clusters and their corresponding annotations are as follows: C0/C4, naïve CD4 T cells; C1/C5, naïve CD8 T cells; C2, regulatory T cells (Tregs); C3-s0/s3, innate-like T/NK-like cells; C3-s1/s2, γδ T cells; C6, central memory-like CD8 T cells; C7, interferon-stimulated gene (ISG)-high naïve CD4 T cells; C8, PLZF⁺ innate-like T cells; C9-s0, activated CD4-like T cells; C9-s1, activated CD8-like T cells; C9-s2, unresolved T cells; C10, effector memory-like CD8 T cells; C11, naïve/memory-like T cells; C12, cycling T cells; C13, PLZF⁺ γδ T cells; and C14, low-quality CD4-like T cells. (B) Dot plot showing the expression of representative marker genes across annotated T cell populations. Dot size represents the percentage of cells expressing each gene within a population, whereas color indicates the scaled average expression level. (C) Tissue-specific UMAP visualizations of T cells from BM, LN, and spleen, displayed using the same UMAP coordinates and cell-type color scheme as in (A). (D) Stacked bar plots showing the relative proportions of annotated T cell populations among the captured T cells within each tissue. Low-quality CD4-like T cells (C14) were excluded, and proportions were recalculated using the remaining cells. (E) Heatmap showing the relative enrichment or depletion of annotated T cell populations across BM, LN, and spleen, quantified using log₂-transformed observed-to-expected (O/E) cell count ratios. Expected counts were calculated from the marginal totals of the cell type-by-tissue contingency table under an independence model. A pseudocount of 0.5 was added to both observed and expected counts before calculating log₂[(O + 0.5)/(E + 0.5)]. Positive and negative values indicate higher- and lower-than-expected representation, respectively, whereas values near zero indicate representation close to expectation. Low-quality CD4-like T cells (C14) were excluded from this analysis.

Although BM, LN, and spleen shared several T cell populations, their relative proportions differed across tissues (Fig. 3C–E). Naïve CD4⁺ T cells accounted for 49.65% of LN T cells and 44.52% of splenic T cells, compared with 7.64% in BM. In contrast, naïve CD8⁺ T cells were the largest annotated population in BM, accounting for 37.69% of T cells. BM also contained higher proportions of innate-like and effector-associated T cell populations. The combined γδ T/NKT-associated population accounted for 16.83% of BM T cells, compared with 1.82% in LN and 3.17% in spleen. PLZF⁺ αβ innate-like T cells represented 9.12% of BM T cells, compared with 0.12% in LN and 3.07% in spleen. KLRG1⁺CX3CR1⁺ effector T cells also accounted for a higher proportion in BM (4.30%) than in LN (0.48%) or spleen (0.93%). These results revealed distinct T cell compositions across the three tissues, with greater representation of innate-like and effector-associated populations in BM.

We next examined transcription factor expression and pySCENIC-inferred regulon activity across T cell populations (Supplementary Fig. S3A,B). Eomes, Klf2, Runx3, and Bach2 associated regulons showed broadly distributed AUCell signals across multiple T cell populations. In contrast, Bhlhe40, Zbtb16, and Maf associated regulons showed distinct activity patterns in innate-like T cell populations. These results revealed both broadly distributed and population associated patterns of inferred regulon activity.

To further examine tissue associated differences within shared T cell populations, we compared mean AUCell scores across BM, LN, and spleen (Supplementary Fig. S3C). Several regulons showed different AUCell scores among tissues within the same T cell population. For example, Ikzf2 associated regulon scores were higher in BM derived Treg cells than in LN and spleen derived Treg cells. These findings suggest that shared T cell populations can exhibit tissue associated differences in inferred regulon activity.

### Regional and compartment-associated heterogeneity of intestinal T cells

To further characterize T cell heterogeneity across intestinal regions, we analyzed intraepithelial lymphocytes (IELs) and lamina propria lymphocytes (LPLs) from the proximal, middle, and distal small intestine and the colon. Eight intestinal samples were included in the analysis (Fig. 4A). After data integration and reclustering, we identified 18 distinct T cell clusters. These clusters included CD8αα-like, CD8αβ, CD4-associated, γδ, regulatory, innate-like, and cycling T cell populations (Fig. 4B). Their identities were supported by the expression of representative marker genes (Fig. 4C and Supplementary Fig. S4).

**Figure 4.**
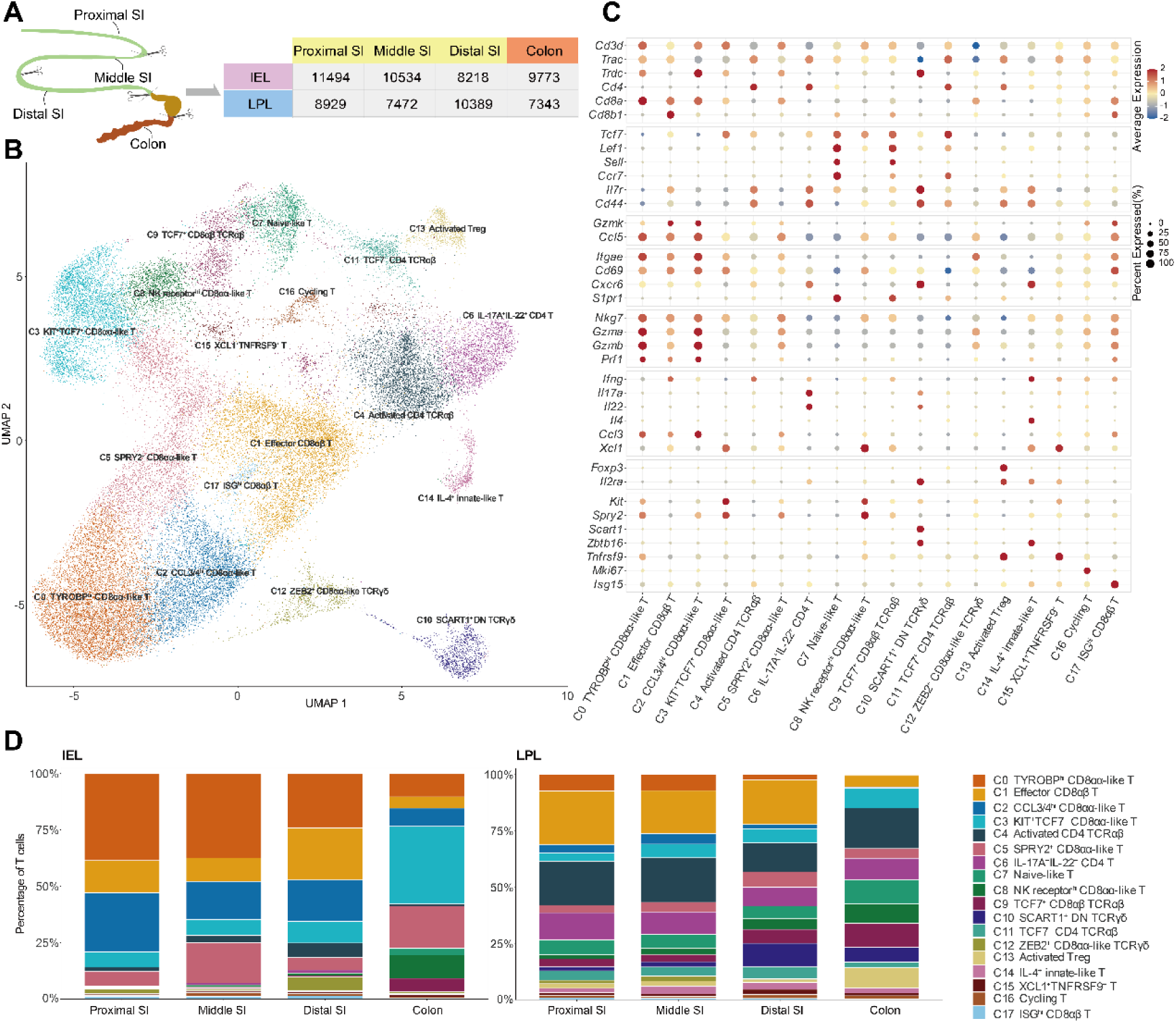
Regional and compartment-associated heterogeneity of intestinal T cells. (A) Schematic illustration of the four intestinal regions, including the proximal, middle, and distal small intestine and the colon. The numbers of T cells recovered from intraepithelial lymphocyte (IEL) and lamina propria lymphocyte (LPL) samples are indicated for each region. SI: small intestine. (B) UMAP visualization of integrated T cells from eight IEL and LPL samples across the four intestinal regions. Each dot represents an individual cell, and colors indicate the 18 annotated T cell clusters. The clusters were annotated as follows: C0, TYROBP-high cytotoxic CD8αα-like TCRαβ/γδ T cells; C1, effector-like CD8αβ TCRαβ TRM-like cells; C2, CCL3/CCL4-high cytotoxic CD8αα-like TCRαβ/γδ T cells; C3, KIT⁺TCF7⁺ CD8αα-like TCRαβ/γδ T cells; C4, activated memory-like CD4⁺ TCRαβ T cells; C5, SPRY2⁺ CD8αα-like TCRαβ/γδ T cells; C6, IL-17A⁺IL-22⁺ CD4⁺ TCRαβ T cells; C7, naive-like T cells; C8, NK receptor-high CD8αα-like TCRαβ/γδ T cells; C9, TCF7⁺ CD8αβ TCRαβ T cells; C10, SCART1⁺ CD4⁻CD8⁻ TCRγδ T cells; C11, TCF7⁺ CD4⁺ TCRαβ T cells; C12, ZEB2⁺ CD8αα-like TCRγδ T cells; C13, activated regulatory T cells; C14, IL-4⁺ innate-like CD4⁺ TCRαβ/γδ T cells; C15, XCL1⁺TNFRSF9⁺ activated CD4/CD8-expressing TCRαβ/γδ T cells; C16, cycling T cells; and C17, ISG-high CD8αβ TCRαβ/γδ T cells. TRM, tissue-resident memory; ISG, interferon-stimulated gene. (C) Dot plot showing the expression of representative lineage- and state-associated marker genes across the annotated intestinal T cell clusters. Dot size indicates the percentage of cells expressing each gene, and color represents the average expression level. (D) Relative proportions of the 18 annotated T cell clusters across the four intestinal regions, shown separately for IEL and LPL populations. Percentages were calculated relative to the total number of T cells within each region-specific IEL or LPL sample.

IELs and LPLs showed distinct T cell compositions (Fig. 4D). Small intestinal IELs contained high proportions of TYROBP-high CD8αα-like T cells (C0), CCL3/4-high CD8αα-like T cells (C2), and effector CD8αβ T cells (C1). In contrast, CD4-associated populations were more abundant in LPLs. These populations included activated CD4 TCRαβ T cells (C4) and IL-17A/IL-22-expressing CD4 T cells (C6).

Different CD8αα-like T cell populations showed different regional distributions and transcriptional features (Supplementary Figs. S4–S5). TYROBP-high CD8αα-like T cells (C0) accounted for 38.68% and 37.55% of proximal and middle small intestinal IELs, respectively. Their proportions decreased to 24.28% in distal small intestinal IELs and 10.43% in colonic IELs. CCL3/4-high CD8αα-like T cells (C2) accounted for 26.32% of proximal small intestinal IELs, compared with 8.08% of colonic IELs. C2 expressed effector differentiation-associated transcription factors, including Zeb2, Runx3, Tbx21, and Prdm1. In contrast, KIT⁺TCF7⁺ CD8αα-like T cells (C3) accounted for 34.39% of colonic IELs, compared with 6.74–9.61% in small intestinal IELs. The expression of Kit and Tcf7 suggested memory-like transcriptional features in this population. NK receptor-high CD8αα-like T cells (C8) also showed higher proportion in colonic IELs (10.32%) than in small intestinal IELs (less than 1% in each region). These results revealed distinct regional distributions of CD8αα-like T cell populations with different transcriptional features.

Regional differences were also observed among LPL populations (Supplementary Fig. S5). SCART1⁺ double-negative γδ T cells (C10) accounted for the highest proportion in distal small intestinal LPLs (10.42%). Activated Treg cells (C13) showed the highest proportion in colonic LPLs (8.93%). The SCART1⁺ γδ T cell population expressed Rorc, Sox13, Zbtb16, and Maf. Activated Treg cells expressed regulatory T cell markers, including Foxp3 and Il2ra (Fig. 4C and Supplementary Fig. S4).

We next used pySCENIC to examine inferred transcription factor regulon activity across intestinal T cell populations (Supplementary Fig. S6). Chd2, Ikzf3, and Zeb1 associated regulons showed broadly distributed AUCell signals across multiple T cell clusters. In contrast, Zbtb16, Foxp3, and Maf associated regulons showed distinct cluster associated activity patterns. We further compared mean AUCell scores across intestinal regions within the same T cell populations (Supplementary Figs. S7–S8). Several regulons showed different AUCell scores among regions within individual T cell clusters. These results revealed both cluster associated and regional variation in inferred regulon activity among intestinal T cell populations.

### Transcriptional diversity and TCR repertoire features of submandibular gland T cells

To further characterize T cell heterogeneity in the submandibular gland (SMG), we analyzed single-cell gene expression and paired TCR V(D)J sequencing data. Transcriptomic analysis identified 13 transcriptional groups, including naïve CD4⁺ and CD8⁺ T cells, central memory-like and EOMES⁺ memory-like CD8⁺ T cells, type I effector-like T cells, helper CD4⁺ T cells, Treg cells, and γδ T17 cells. Interferon-responsive and stress-associated T cell populations were also identified (Fig. 5A). These populations were distinguished by the expression of representative marker genes (Fig. 5B).

**Figure 5.**
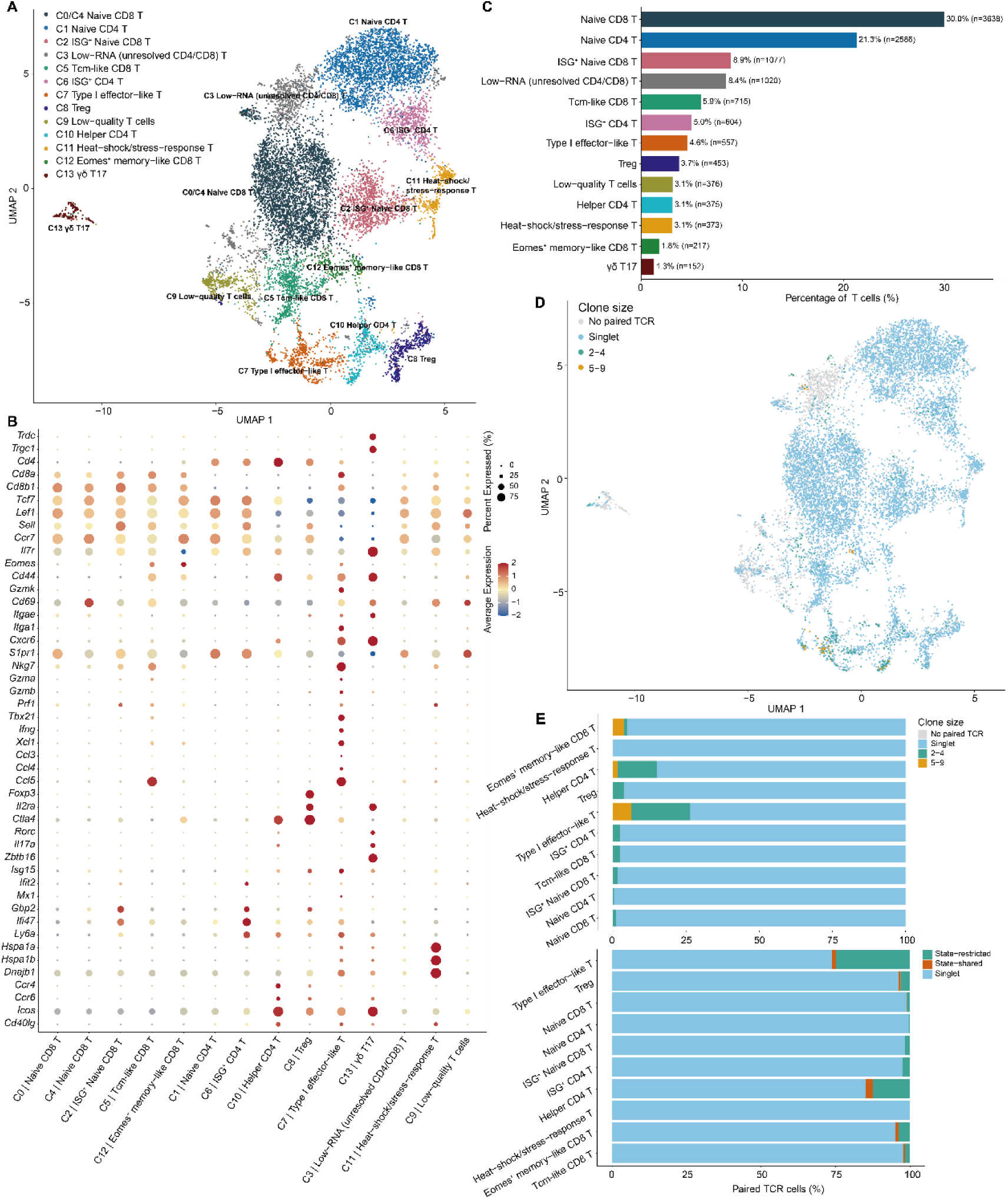
Single-cell transcriptomic and TCR repertoire characterization of T cells from submandibular glands. (A) UMAP visualization of T cells isolated from the submandibular glands (SMG) of healthy mice, showing transcriptionally defined T cell clusters annotated according to canonical lineage and functional marker genes. Each dot represents an individual cell, and colors indicate annotated T cell populations. (B) Dot plot showing the expression of representative T cell lineage, differentiation, activation, effector, memory, and tissue-residency-associated genes across the annotated populations. Dot size represents the percentage of cells expressing each gene, and color indicates scaled average expression. (C) Relative abundance of the annotated T cell populations in the SMG dataset. Bars indicate the percentage of total T cells represented by each population, with the corresponding cell numbers (n) shown. (D) UMAP visualization of TCR clonotype sizes in SMG T cells with paired TCR sequencing data. Cells are colored by clonotype size: Singlet (one cell), 2–4 cells, or ≥5 cells. (E) Distribution of clonotype sizes and sharing patterns across selected T cell populations with paired TCR sequences. The upper panel shows the proportions of paired-TCR cells assigned to Singlet (one cell per clonotype) or expanded clonotypes, while the lower panel shows the proportions assigned to Singlet, state-restricted, or state-shared clonotypes. State-restricted clonotypes were detected within a single annotated T cell population, whereas state-shared clonotypes were detected across multiple populations. The low-RNA (C3), low-quality (C9), and γδ T17 (C13) populations were excluded from this analysis. αβ TCR clonotypes were defined by identical paired TRA and TRB CDR3 nucleotide sequences.

Naïve CD8⁺ T cells were the largest population, accounting for 30.0% of SMG T cells (n = 3,638), followed by naïve CD4⁺ T cells (21.3%, n = 2,586). Other populations included ISG⁺ naïve CD8⁺ T cells (8.9%), central memory-like CD8⁺ T cells (5.9%), ISG⁺ CD4⁺ T cells (5.0%), and type I effector-like T cells (4.6%) (Fig. 5C). Compared with the BM, LN, and spleen populations analyzed in Fig. 3, SMG shared several major T cell states but showed a different cellular composition. A distinct population of type I effector-like T cells was identified in SMG. In contrast to the BM, LN, and spleen analysis, no distinct cycling T cell cluster or PLZF⁺ innate-like T cell cluster was identified in SMG. These findings revealed differences in the relative abundance and transcriptional states of T cell populations across these tissues.

We next examined the expression of tissue residency-associated genes (Fig. 5B). *Cxcr6* expression was detected in type I effector-like and γδ T17 cells, whereas *Itga1* and *Itgae* showed limited expression across most T cell populations. Although SMG tissues were perfused before collection, the recovered cells included naïve, memory like, and effector associated transcriptional states.

We next examined TCR clonotypes across the identified T cell populations (Fig. 5D-E). Most cells with assigned TCR clonotypes belonged to singleton clonotypes, whereas a smaller proportion belonged to clonotypes shared by multiple cells. The distribution of expanded clonotypes differed across T cell populations. Type I effector-like T cells contained a higher proportion of cells belonging to expanded clonotypes. Most recurrent clonotypes were detected within a single transcriptional state, although six expanded αβ TCR clonotypes were detected across multiple states (data not shown). These findings revealed population-associated differences in TCR clonal expansion, with greater representation of expanded clonotypes among type I effector-like T cells.

## Discussion

In this study, we constructed an integrated single-cell transcriptional atlas of T cells from multiple lymphoid and non-lymphoid tissues of healthy mice. We identified shared T cell states with distinct cellular compositions and transcriptional features across organs. These findings indicate that tissue-associated T cell heterogeneity involves not only differences in population abundance but also differences in differentiation and function associated transcriptional states. Further analyses revealed diverse memory-like, innate-like, and effector associated T cell populations in bone marrow, regional heterogeneity of intestinal CD8αα-like T cells, and distinct transcriptional states in the submandibular gland. Together, these findings provide a reference for understanding T cell diversity across healthy tissues.

BM and SMG showed different T cell compositions compared with LN and spleen. BM showed higher proportion of innate-like T cells and contained distinct effector memory-like T cell populations. Previous studies have identified CD4⁻CD8⁻ TCRαβ⁺ innate-like T cells that accumulate in the BM of healthy mice and exhibit potent innate immune functions^22^. In addition, γδ T cells and NKT-like T cells may contribute to rapid immune responses through their innate-like effector functions. These findings suggest that BM may support diverse innate-like T cell populations involved in immune surveillance. Meanwhile, BM also serves as an important reservoir for memory T cells. Studies in mice and humans have shown that BM provides specialized survival niches for memory T cells^23, 24^, allowing them to persist over time while retaining the capacity for strong recall responses upon antigen re-exposure. These findings suggest that BM may support both rapid immune surveillance by innate-like T cells and long-term immunological memory maintained by memory T cells.

In contrast, SMG contained distinct cell populations of Helper CD4 T cells, ISG⁺ CD4 T cells, and Type I effector-like T cells. Previous studies have shown that salivary glands can support local helper T cell responses, including Th1 associated responses, and that effector CD4 T cells contribute to antiviral defense in the salivary gland^25, 26^. These findings suggest that helper and effector associated T cells may play important roles in local immunity. In our study, the presence of these populations in healthy SMG indicates diverse T cell transcriptional states even in the absence of overt inflammation. However, the functional roles and origins of these populations remain unclear. Further studies are needed to determine whether these transcriptional states are established before T cells enter the SMG or are shaped by the local salivary gland environment.

Intestinal T cells showed heterogeneity between the intraepithelial and lamina propria compartments and across intestinal regions. Previous studies have demonstrated that conventional and regulatory CD4⁺ T cells can undergo progressive transcriptional reprogramming as they migrate from the lamina propria into the intestinal epithelium^27, 28^. This process involves downregulation of ThPOK, increased *Runx3* expression, and acquisition of an IEL-associated transcriptional program. These findings highlight the role of the intestinal epithelium in shaping T cell states. In our study, distinct CD8αα-like T cell populations also showed different regional distributions. TYROBP-high and CCL3/4-high populations were more abundant in proximal small intestinal IELs, whereas KIT⁺TCF7⁺ cells were more prominent in colonic IELs. Recent studies have similarly identified Tcf7⁺ memory-like and Prdm1⁺ effector-like states among intestinal CD8αα IELs, with different distributions between the small and large intestine^29^. Together, these observations suggest that both tissue compartment and intestinal region are associated with T cell transcriptional heterogeneity. However, the developmental origins of the CD8αα-like populations identified here and their relationships to LPL populations remain unclear.

Several limitations should be considered. Cells from multiple mice were pooled for each tissue sample without independent biological replicates. Therefore, between animal variation could not be assessed, and tissue associated differences could not be evaluated using biological replicate level statistical tests. Differences in tissue processing, sequencing platforms, and potential contributions from circulating T cells may also influence the observed cellular compositions and transcriptional patterns. In addition, tissue specific reclustering results depend on cell numbers and clustering resolution. The absence of a distinct cluster in one tissue does not necessarily indicate the absence of the corresponding T cell state. Pseudotime trajectories and pySCENIC regulon activities are computational inferences and do not directly establish developmental relationships or transcription factor function. Further studies with independent biological replicates and functional validation will be needed to determine the biological significance of these findings. Overall, this study provides a single-cell reference for investigating shared and tissue associated T cell heterogeneity across multiple organs of healthy mice.

## Materials and methods

### Tissue dissociation and preparation of single-cell suspensions

Spleen, thymus, and lymph nodes were harvested from healthy mice and mechanically dissociated in ice-cold phosphate-buffered saline (PBS) supplemented with 2% fetal bovine serum (FBS). The resulting cell suspensions were passed through 70-μm cell strainers. Splenic and thymic cells were treated with red blood cell (RBC) lysis buffer for 5 min at room temperature, whereas lymph node cells were processed without routine RBC lysis.

Bone marrow cells were isolated by flushing the femurs and tibias with PBS supplemented with 2% FBS. The resulting cell suspensions were filtered through 70-μm cell strainers, followed by RBC lysis for 10 min at room temperature.

Intraepithelial lymphocytes (IELs) and lamina propria lymphocytes (LPLs) were isolated separately from the small intestine and colon. Following removal of surrounding adipose tissue and Peyer’s patches, intestinal segments were opened longitudinally and thoroughly washed with PBS to remove luminal contents.

For IEL isolation, intestinal tissues were incubated in RPMI 1640 medium supplemented with approximately 1 mM dithiothreitol (DTT), 5 mM EDTA, and 2% FBS at 37°C with agitation for 20 min (small intestine) or 25 min (colon). The released cells were collected and filtered.

For LPL isolation, the remaining intestinal tissues were washed, finely minced, and enzymatically digested in RPMI 1640 medium containing 0.5 mg/mL dispase II, 1 mg/mL collagenase type II, 10 μg/mL DNase I, and 2% FBS at 37°C for 45 min (small intestine) or 60 min (colon). Following digestion, the resulting cell suspensions were filtered.

Submandibular glands were dissected from mice, minced, and enzymatically digested in RPMI 1640 medium supplemented with 1 mg/mL collagenase type II, 10 μg/mL DNase I, and 2% FBS at 37°C for 30 min. The resulting cell suspensions were passed through 70-μm cell strainers.

Cell suspensions were collected by centrifugation at 400 × g for 5 min at 4°C, except for intestinal cell suspensions, which were centrifuged at 500 × g for 10 min. Intestinal IELs, LPLs, and submandibular gland-derived lymphocytes were enriched using discontinuous 40%/80% Percoll density gradients. Density-gradient centrifugation was performed at 2,500 rpm for 20 min at 4°C, with acceleration and deceleration settings of 5 and 1, respectively. Lymphocytes were collected and resuspended in fluorescence-activated cell sorting (FACS) buffer for subsequent antibody staining and flow cytometric sorting.

### Flow cytometry and T cell sorting

Single-cell suspensions obtained from different tissues were incubated with TruStain FcX™ (anti-mouse CD16/32, BioLegend) on ice for 10 min to block nonspecific Fc receptor-mediated antibody binding. Cells were subsequently stained with eFluor 780 fixable viability dye (1:1,000; Thermo Fisher Scientific) and FITC anti-mouse CD3 antibody (1:100; BioLegend). Following staining, single, viable CD3 T cells were isolated using a Sony MA900 cell sorter or a BD FACS Melody cell sorter. The sorted CD3 T cells were collected for subsequent single-cell RNA sequencing (scRNA-seq).

### Single-cell RNA sequencing and TCR V(D)J library preparation

Sorted CD3⁺ T cells from all tissues except the submandibular glands were subjected to 3’ single-cell RNA sequencing using the MobiNova platform.

Freshly prepared single-cell suspensions were adjusted to a concentration of 700-1,200 cells/μL. Single-cell capture and library preparation were performed using the MobiCube High-throughput Single Cell 3’ Transcriptome Kit V2.5 (catalog no. S050200601) on the MobiNova-100 microfluidic platform according to the manufacturer’s instructions. The libraries were sequenced on an Illumina NovaSeq X Plus platform with 150-bp paired-end reads (PE150).

Sorted CD3⁺ T cells from the submandibular glands were subjected to 5’ single-cell gene expression profiling and T cell receptor (TCR) V(D)J sequencing using the 10x Genomics Chromium platform. Gene expression libraries were prepared using the Chromium Next GEM Single Cell 5′ Kit v3 (10x Genomics, PN-1000699), according to the manufacturer’s instructions. TCR V(D)J libraries were prepared using the Chromium Single Cell Mouse TCR Amplification Kit (PN-1000254). The constructed libraries were sequenced on the BGI DNBSEQ-T7-PE150 platform for high-throughput sequencing.

### Processing of raw single-cell sequencing data

Raw sequencing data generated using the MobiNova platform were processed using MobiVision software (version 3.2) with default parameters. Sequencing reads were aligned to the mouse reference genome (GRCm39), and cell barcode identification, cell filtering, and gene expression quantification were performed using the software pipeline. The resulting filtered feature–barcode matrices, containing unique molecular identifier (UMI) counts for individual genes and cells, were used for downstream analyses.

Raw gene expression and TCR V(D)J sequencing data from submandibular gland-derived T cells were processed jointly using Cell Ranger (version 9.0.1; 10x Genomics) with the ‘cellranger mult’ pipeline. Gene expression reads were aligned to the mouse reference transcriptome (refdata-gex-GRCm39-2024-A), whereas TCR V(D)J reads were processed using the mouse V(D)J reference (refdata-cellranger-vdj-GRCm38-alts-ensembl-7.0.0). Gene expression and V(D)J libraries were specified separately in the multi-library configuration, with the expected number of recovered cells set to 1,000.

The Cell Ranger pipeline was used to generate gene expression count matrices and assemble TCR V(D)J sequences. The resulting transcriptomic and TCR data were linked using their corresponding cell barcodes to enable joint analysis of T cell transcriptional states and receptor repertoires.

### Quality control and downstream analysis of single-cell RNA-seq data

Single-cell gene expression matrices from 13 samples were imported into Seurat (version 5.3.0)^30^ for quality control and downstream analysis. Cells were retained if they expressed more than 500 and fewer than 6,500 genes (nFeature_RNA), contained fewer than 50,000 UMIs (nCount_RNA), and had a mitochondrial transcript fraction below 15%.

Potential doublets were identified independently for each sample using DoubletFinder (version 2.0.6)^31^. For doublet detection, gene expression data were normalized using the LogNormalize method with a scaling factor of 10,000. The top 2,000 highly variable genes were identified using the variance-stabilizing transformation (vst) method, followed by data scaling and principal component analysis (PCA).

DoubletFinder was performed using the first 30 principal components, with pN set to 0.25. The optimal pK value was determined separately for each sample by maximizing the mean–variance-normalized bimodality coefficient (BCmvn). The expected doublet rate was set to 5% of the number of cells remaining after initial quality control, without homotypic doublet adjustment. Cells classified as doublets were excluded, and the remaining singlets were retained for subsequent analyses.

### Data integration, dimensionality reduction, and clustering

Following quality control and doublet removal, T cells from all 13 samples were merged into a single Seurat object to construct the multi-tissue T cell atlas. Gene expression data were normalized using the LogNormalize method with a scaling factor of 10,000. The top 5,000 highly variable genes were identified using the vst method, and their expression values were scaled before PCA. A total of 50 principal components were calculated.

To account for sample-associated batch effects, Harmony (version 1.2.3)^32^ was applied to the first 30 principal components using sample identity as the batch-correction variable. UMAP embeddings were generated both before and after Harmony correction to assess sample-associated differences. The Harmony-corrected embeddings were subsequently used for downstream dimensionality reduction, graph construction, and clustering.

A nearest-neighbor graph was constructed using the first 30 Harmony dimensions. Unsupervised graph-based clustering was performed using the Louvain algorithm (algorithm = 1) with a resolution of 0.6. UMAP visualization was generated using the uwot implementation with cosine distance.

### Identification of cluster-specific marker genes and T cell annotation

Cluster-specific marker genes were identified using the FindAllMarkers function in Seurat after joining the RNA assay layers. Differential expression testing was performed using the Wilcoxon rank-sum test, retaining positively enriched genes expressed in at least 20% of cells in either comparison group (min.pct = 0.20) and applying a minimum log2 fold-change threshold of 0.25.

Significant cluster-specific markers were further defined using an adjusted P value below 0.05 and an average log2 fold change greater than 0.5. T cell populations were characterized based on their transcriptional signatures and the expression of established T cell lineage and functional markers.

### Transcription factor regulon inference and activity analysis

Transcription factor (TF) regulatory networks were inferred using pySCENIC (version 0.12.1)^33, 34^. Briefly, GRNBoost2^35^ was used to identify candidate TF– target gene interactions, followed by cisTarget motif enrichment analysis to identify motif-supported regulons. Regulon activity was then calculated for individual cells using AUCell.

Analyses were performed independently for the integrated atlas and tissue-specific datasets. Regulon activity patterns were examined across T-cell populations and along thymocyte pseudotime trajectories. Mean AUCell scores were further compared within shared T-cell populations across BM, LN, and spleen, and within intestinal T-cell populations across regions and IEL/LPL compartments. Only population–tissue or population–region groups containing at least 50 cells were included in these comparisons.

### Pseudotime trajectory inference using Slingshot

Pseudotime analysis of thymic T cells was performed using Slingshot (version 2.14.0)^36^. NKT-like and proliferating DP populations were excluded, and the remaining clusters were analyzed using the first 30 principal components. The early DP population was designated as the starting cluster. Slingshot was used to infer maturation-associated trajectories, and the resulting pseudotime values were visualized alongside the expression of representative thymocyte developmental and maturation-associated genes.

### T cell receptor repertoire and clonotype analysis

T cell receptor (TCR) repertoire analysis was performed using paired single-cell V(D)J sequencing data from submandibular gland T cells. Cells with productive, high-confidence paired TRA and TRB sequences were retained for conventional αβ T cell repertoire analysis. Clonotypes were defined based on identical paired TRA and TRB CDR3 sequences, and clonotypes detected in two or more cells were classified as recurrent clonotypes. Clonotype distribution and recurrence were subsequently examined across annotated T cell populations.

## Data availability

The raw sequencing data generated in this study, including MobiNova-based 3’ single-cell RNA sequencing (scRNA-seq) data and 10x Genomics-based 5’ scRNA-seq and T cell receptor (TCR) V(D)J sequencing data, have been deposited in the Genome Sequence Archive^37^ in National Genomics Data Center^38^, China National Center for Bioinformation/Beijing Institute of Genomics, Chinese Academy of Sciences (GSA: CRA053394) that are publicly accessible at https://ngdc.cncb.ac.cn/gsa.

## Author contributions

Xiaojing.W. and X.Z. conceived the project. Xiaojing.W. wrote the manuscript. Xiaojing.W. performed isolation of thymus, bone marrow, spleen, lymph node, intestine, and single-cell sequencing data analysis. B.P. performed isolation of submandibular gland and single-cell sequencing. J.L., Z.H., Xueqing.W., Y.W. performed mouse dissection and organs isolation. X.Z. obtained the funding and supervised the research.

## Conflict of interest

The authors declare no conflict of interests.

## Acknowledgement

X.Z. is supported by Innovative Drug Research and Development National Science and Technology Major Project (2026ZD1805700), the Strategic Priority Research Program of the Chinese Academy of Science (grant No. XDB0990000).

---

**Supplementary Fig. S1.**
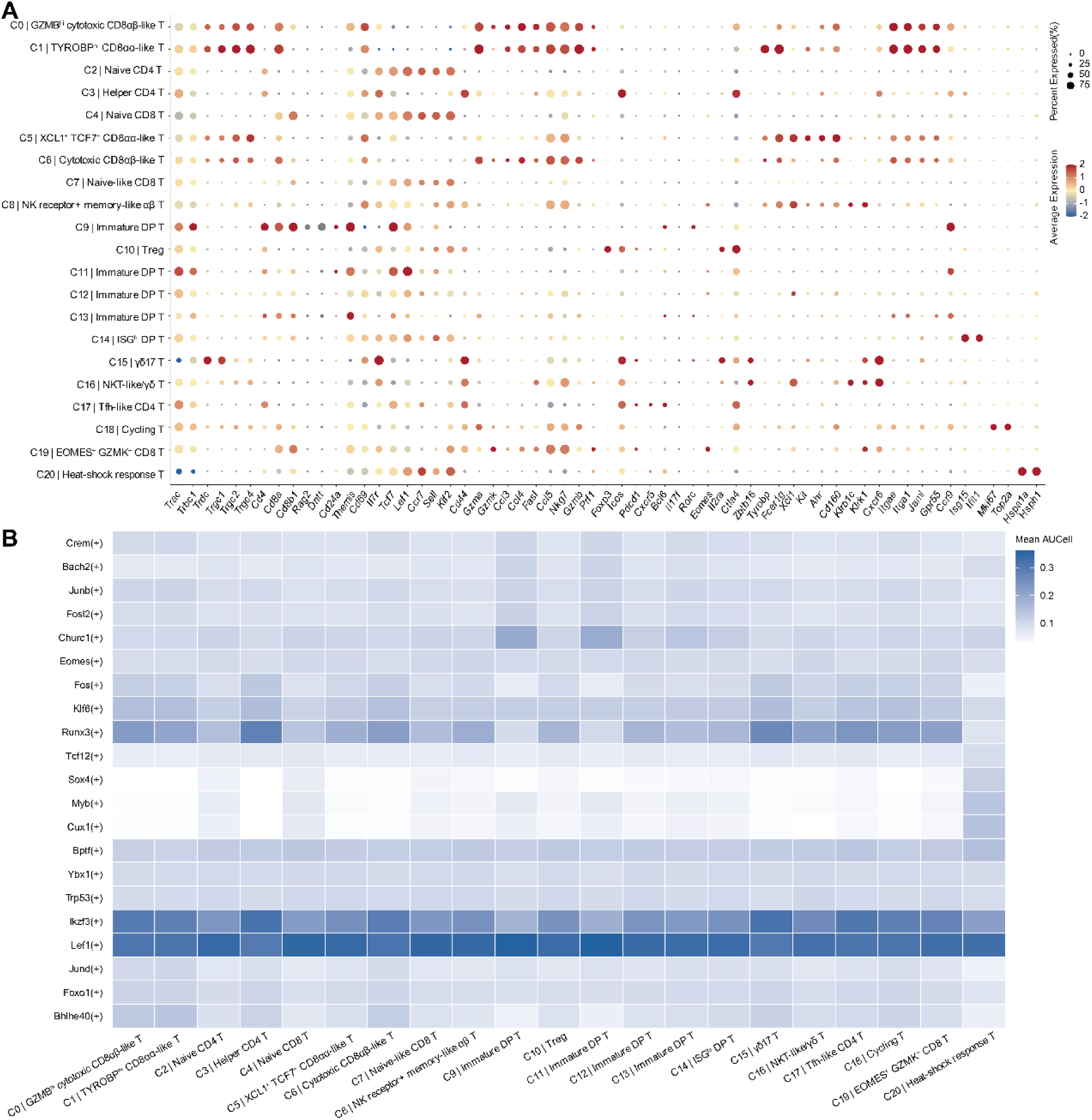
Marker gene expression and inferred regulon activity in the integrated T cell atlas. (A) Marker gene expression profiles of T cell populations across tissues. Dot plot showing the expression patterns of representative marker genes across the annotated T cell clusters identified in the integrated single-cell RNA-sequencing dataset. Dot size indicates the percentage of cells expressing each gene within a cluster, whereas color represents the scaled average expression level. (B) Heatmap showing the mean AUCell scores of pySCENIC-inferred regulons across T cell populations. The top 10 regulons ranked by mean AUCell score were selected independently for each cluster, and their union was used to construct the heatmap. Duplicate regulons were included only once. Color intensity represents the mean AUCell score for each regulon within each T cell population, with darker blue indicating higher scores.

**Supplementary Fig. S2.**
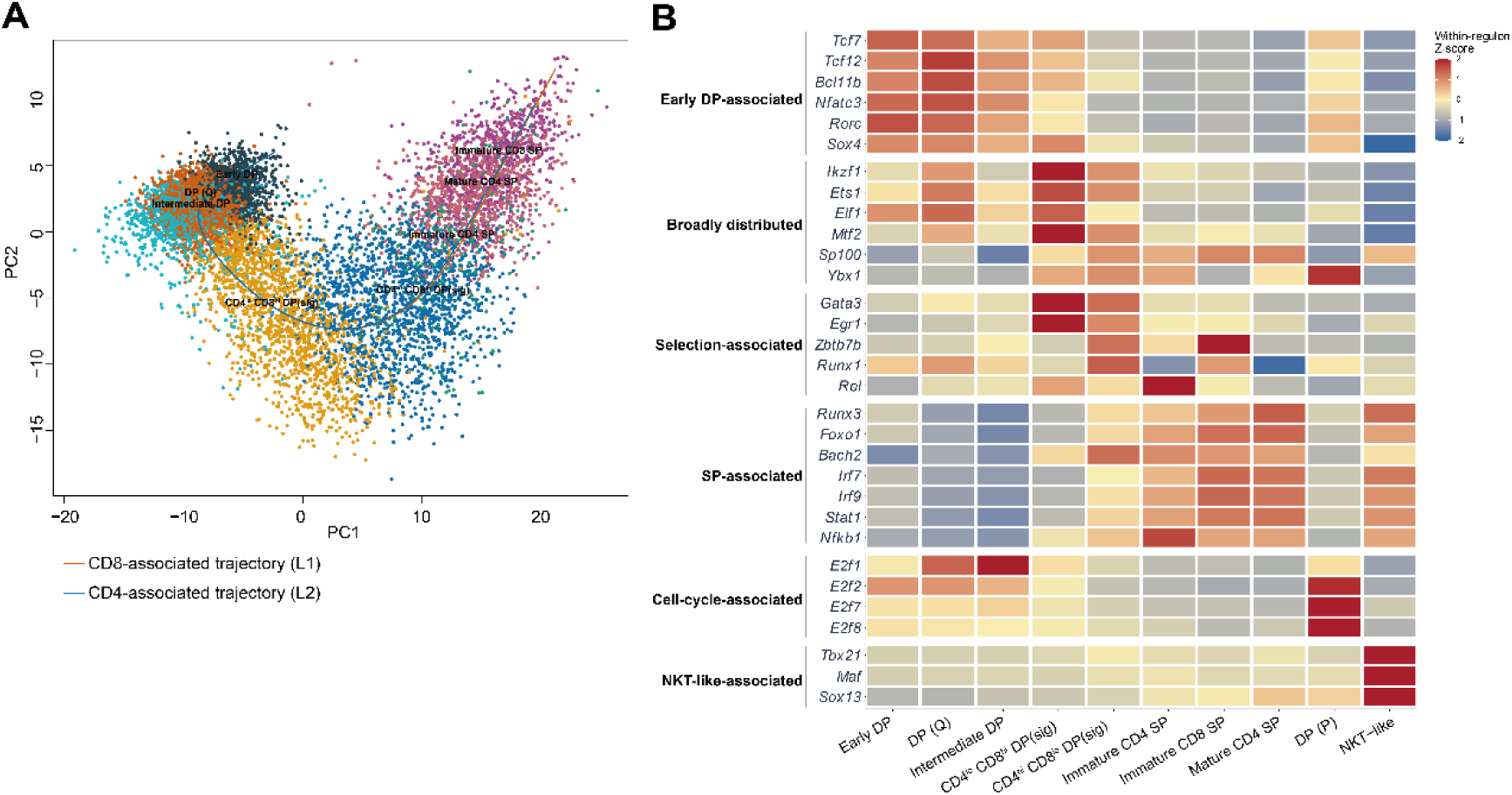
Developmental trajectories and transcriptional regulatory programs of thymic T cells. (A) Principal component analysis (PCA) of thymic T cells showing distinct developmental populations. Slingshot-inferred trajectories are overlaid to illustrate CD8-associated (L1, orange) and CD4-associated (L2, blue) developmental paths originating from the early DP population. (B) Heatmap showing the relative activity of selected transcription factor regulons across thymocyte developmental states, inferred using pySCENIC. Regulons are grouped into putative functional categories, including early DP-enriched, broadly distributed, selection/transition-associated, SP-associated/maturation, proliferating DP-associated, and NKT-like-associated programs. Colors represent within-regulon Z-scores calculated from the mean AUCell scores across cell clusters, with red and blue indicating relatively higher and lower regulon activity, respectively.

**Supplementary Fig. S3.**
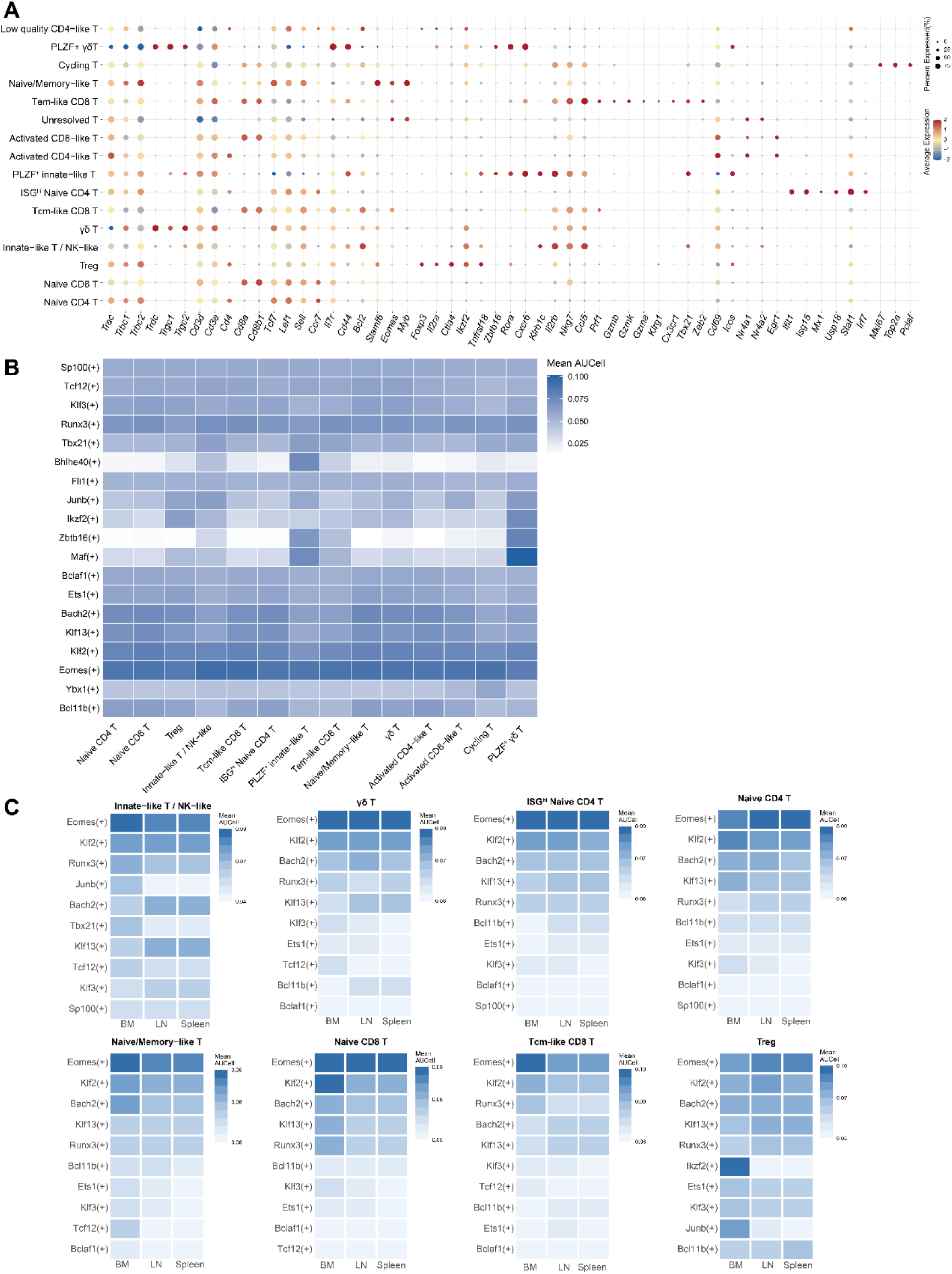
Marker gene expression and inferred regulon activity across bone marrow, lymph node, and spleen T cell populations. (A) Dot plot showing the expression of representative marker genes across T cell clusters identified from the bone marrow, lymph node, and spleen single-cell RNA-sequencing datasets. Dot size represents the percentage of cells expressing each gene within a cluster, while color indicates the scaled average expression level. (B) Heatmap showing the mean AUCell scores of pySCENIC-inferred regulons across T cell populations. The top 10 regulons ranked by mean AUCell score were identified independently for each population, and their union was used for visualization. Color intensity indicates the mean AUCell score. (C) Heatmaps showing tissue-specific mean AUCell scores for the top 10 regulons of the indicated T cell populations across bone marrow (BM), lymph node (LN), and spleen. Regulons were selected based on their mean AUCell scores across all cells of the corresponding population, independently of tissue origin. Only population-tissue combinations containing at least 50 cells were included. Each population was displayed using an independent color scale to highlight within-population differences across tissues.

**Supplementary Fig. S4.**
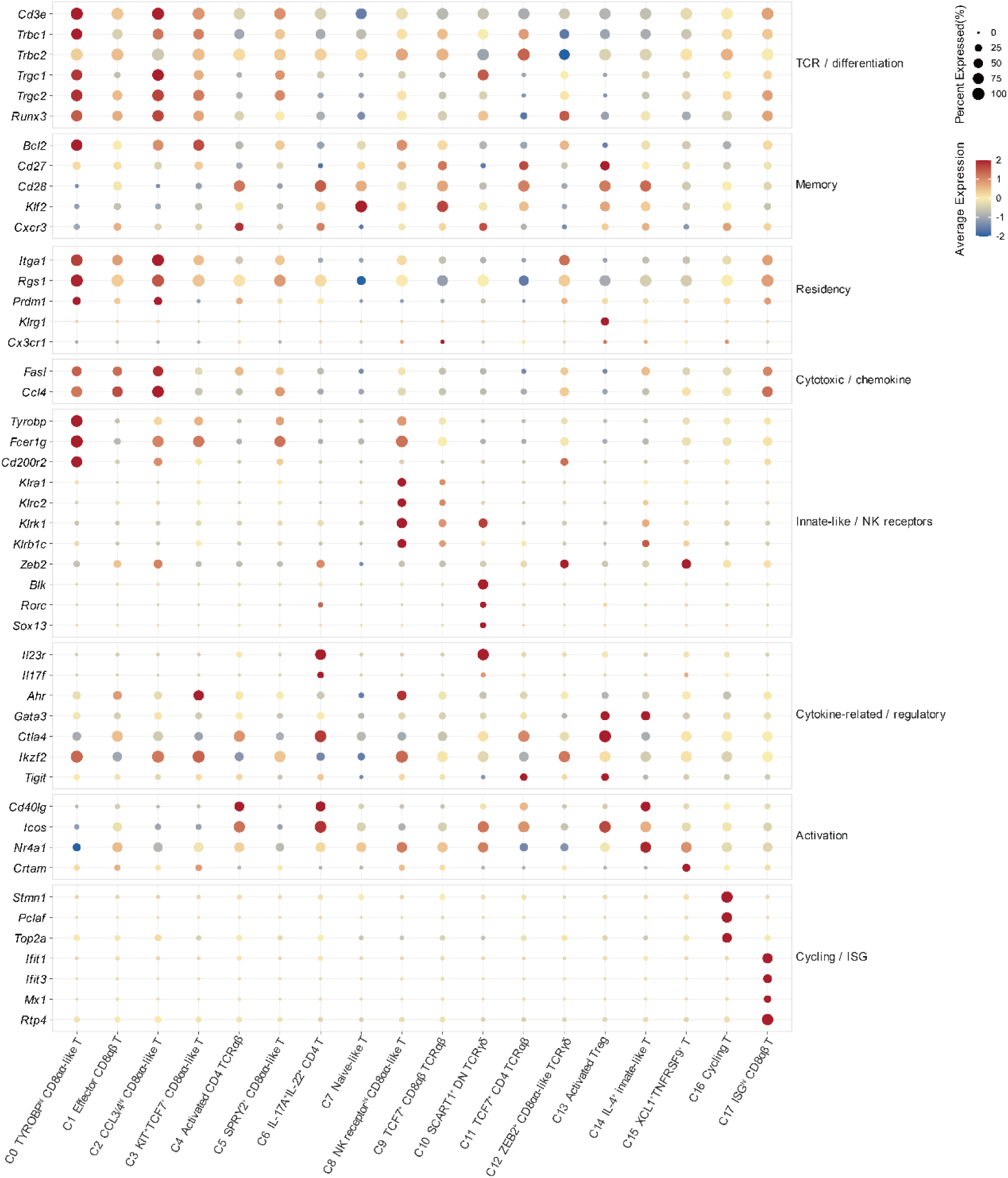
Transcriptional characterization of intestinal T cell subsets based on marker gene expression. Dot plots showing the expression patterns of representative functional marker genes across intestinal T cell subpopulations. Dot size represents the percentage of cells expressing each gene within a cluster, and color indicates the scaled average expression level.

**Supplementary Fig. S5.**
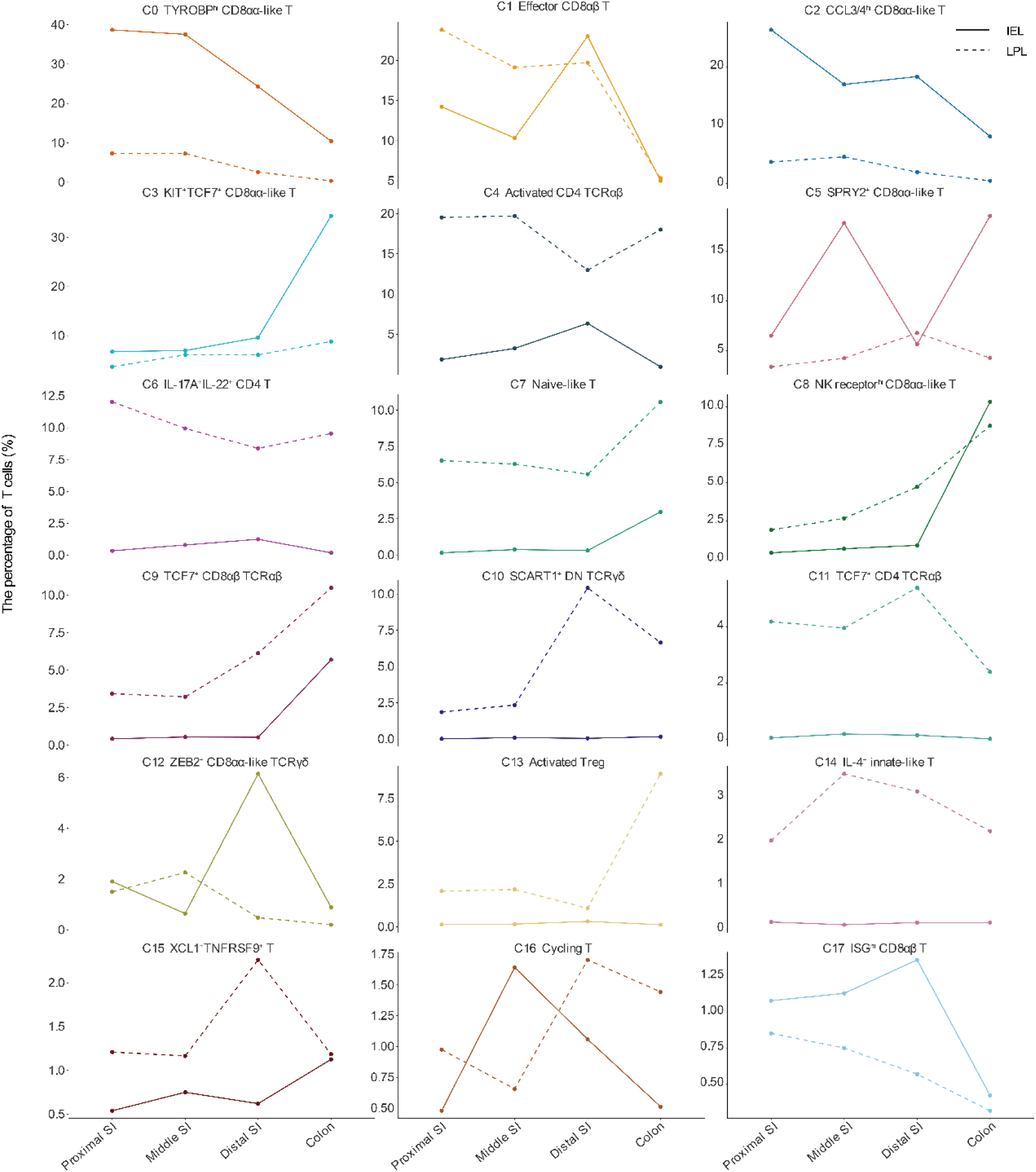
Regional distribution of intestinal T cell populations across intestinal compartments. Line plots showing the relative abundance of annotated intestinal T cell subpopulations across the proximal small intestine (Proximal SI), middle small intestine (Middle SI), distal small intestine (Distal SI), and colon. Each panel represents an individual T cell cluster, with colors distinguishing the annotated populations. Solid and dashed lines indicate intraepithelial lymphocytes (IEL) and lamina propria lymphocytes (LPL), respectively. The y-axis represents the percentage of T cells within the corresponding intestinal compartment, and the x-axis indicates the intestinal region.

**Supplementary Fig. S6.**
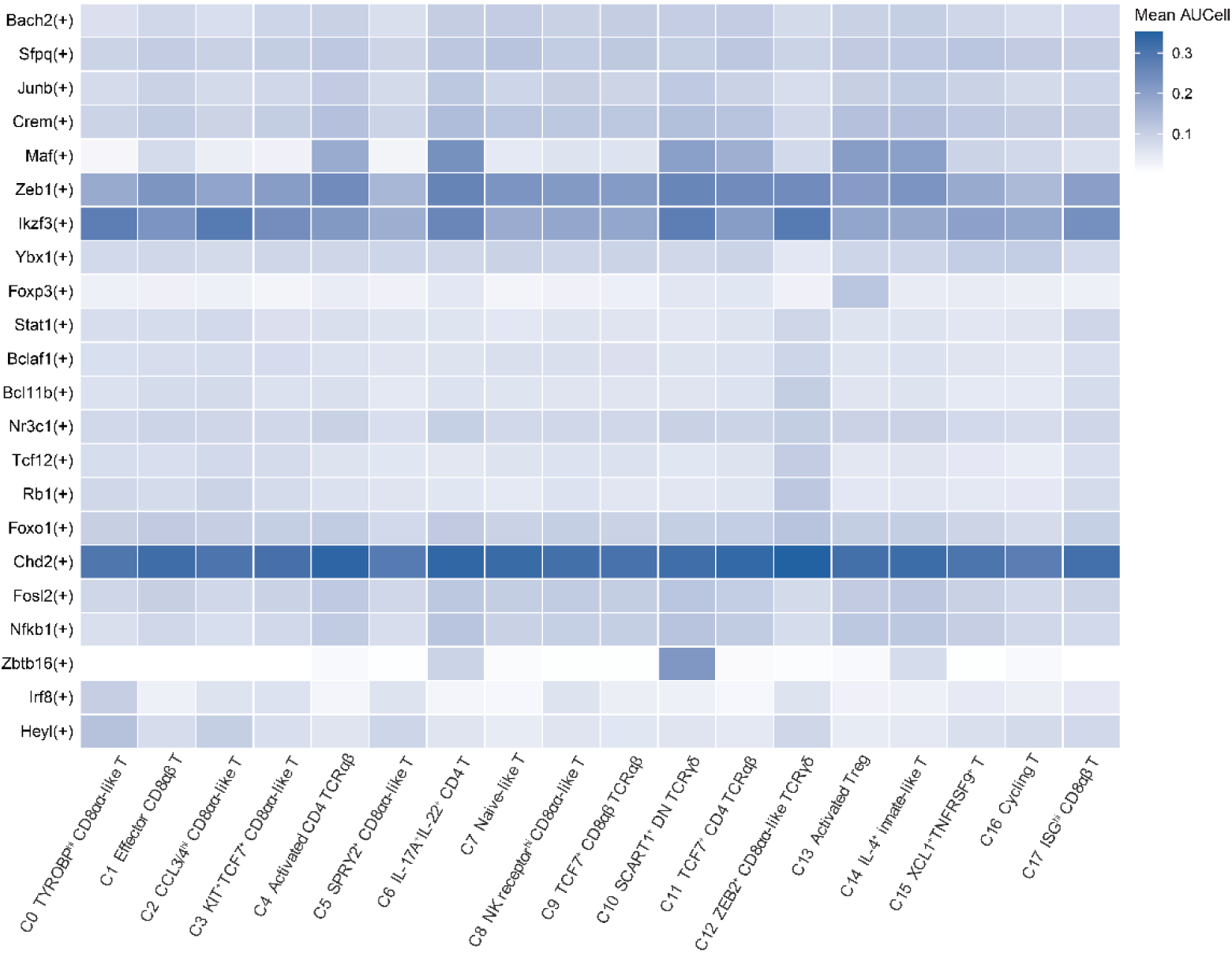
Distribution of inferred regulon activity across intestinal T cell clusters. Heatmap showing the mean AUCell scores of pySCENIC-inferred regulons across 18 intestinal T cell clusters (C0-C17). For each cluster, the top 10 regulons were selected based on their mean AUCell scores. The union of these regulons, with duplicates removed, was used to construct the heatmap. Color intensity indicates the mean AUCell score, with darker blue representing higher scores.

**Supplementary Fig. S7.**
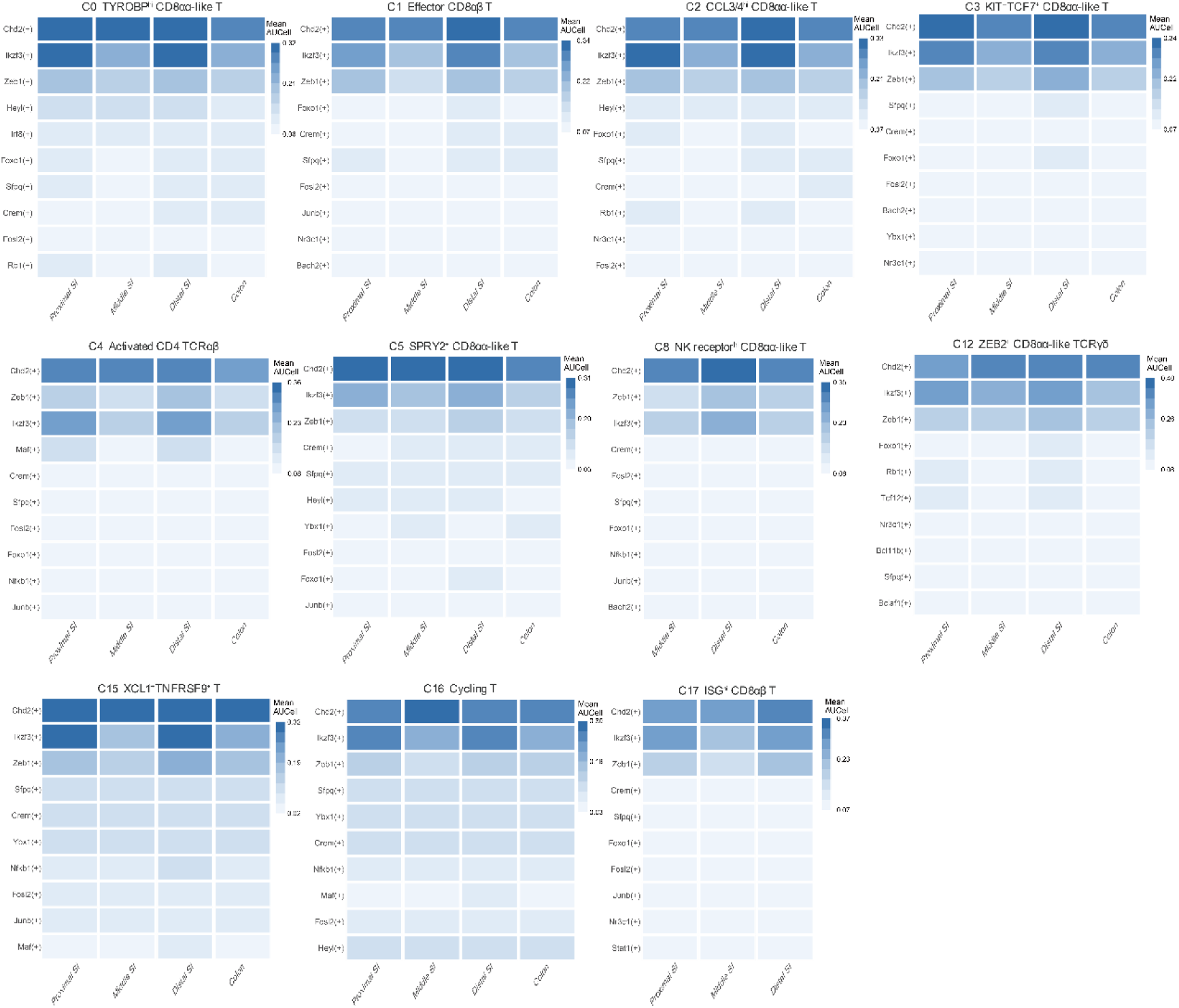
Regional variation in inferred regulon activity among intestinal intraepithelial lymphocytes. Heatmaps showing the mean AUCell scores of the top 10 pySCENIC-inferred regulons for individual T cell clusters within the intestinal intraepithelial lymphocyte (IEL) compartment. For each cluster, regulons were selected according to their mean AUCell scores across all intestinal cells belonging to that cluster, independently of anatomical region or compartment. Columns represent the proximal small intestine (Proximal SI), middle small intestine (Middle SI), distal small intestine (Distal SI), and colon, while rows represent the selected regulons. Only cluster-region combinations containing at least 50 IEL cells were included.

**Supplementary Fig. S8.**
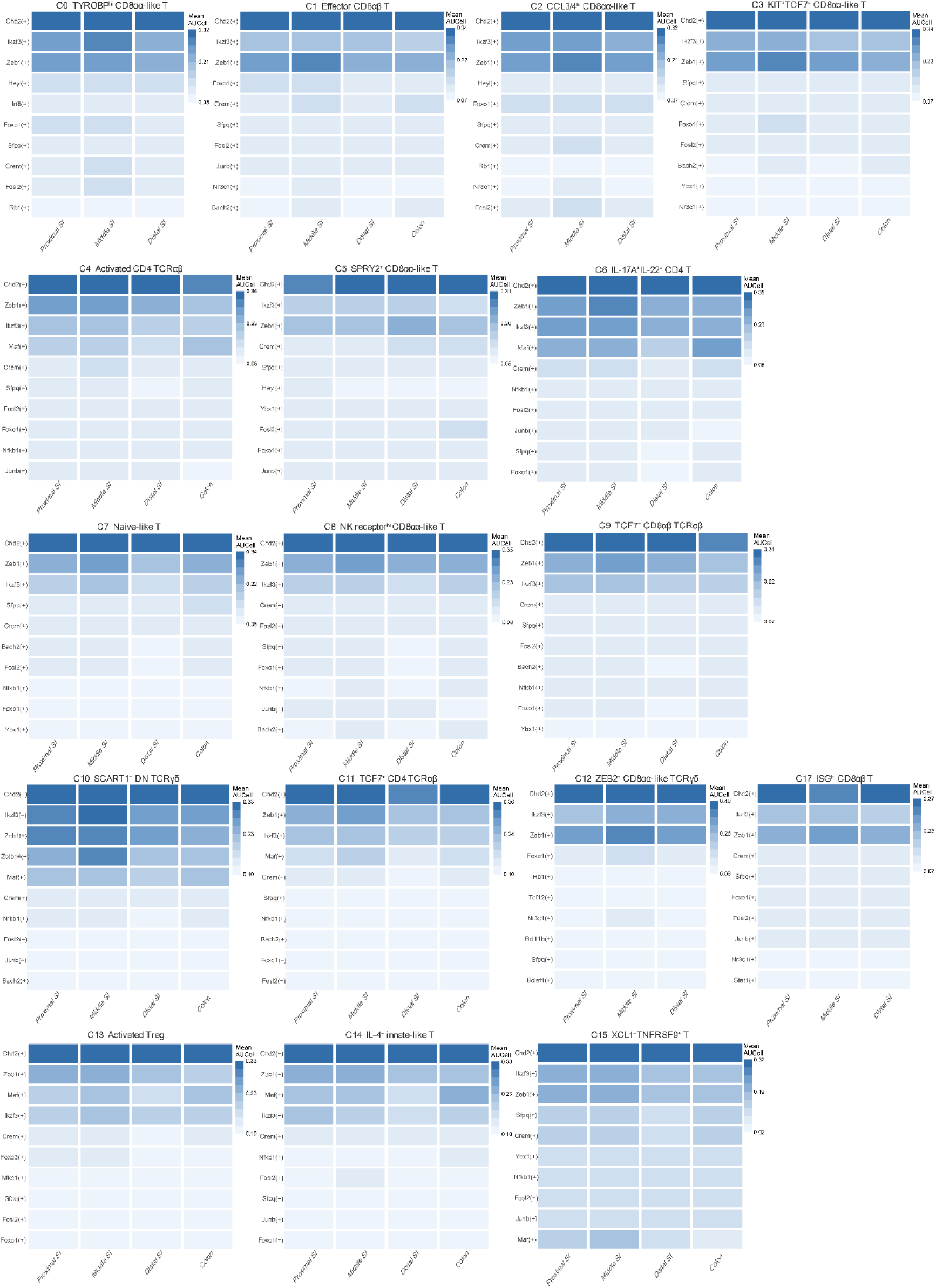
Regional variation in inferred regulon activity among intestinal lamina propria lymphocytes. Heatmaps showing the mean AUCell scores of the top 10 pySCENIC-inferred regulons for individual T cell clusters within the intestinal lamina propria lymphocyte (LPL) compartment. Only cluster-region combinations containing at least 50 LPL cells were included.

## Notes

### Competing Interest Statement

The authors have declared no competing interest.

